# An approach to enhance symbiotic nitrogen fixation

**DOI:** 10.1101/2025.04.23.649995

**Authors:** Aleksandr Gavrin, Marcelo Bueno Batista, Edouard Evangelisti, Ray Dixon, Sebastian Schornack

## Abstract

Through symbiosis with nitrogen-fixing bacteria, cultivated legumes provide themselves and subsequent crops with nitrogen, making genetic improvements of symbiotic efficiency particularly attractive. Here, we identify the symbiosis-specific *GBP1* gene which negatively regulates nitrogen fixation by attenuating bacterial nitrogenase activity in *Medicago truncatula* nodules. *GBP1* inactivation increases nitrogen fixation without affecting nodule development and numbers, providing inroads for engineering legumes with increased productivity for sustainable nitrogen provision.

## Main

Biological nitrogen fixation is the primary source of plant-available nitrogen in many ecosystems [1]. The *Rhizobium*-legume symbiosis is one of the most productive nitrogen-fixing systems. In this so-called root nodule symbiosis, bacteria are accommodated within plant cells of special organs, the root nodules, where they bind elementary nitrogen from the air. As a result of this symbiosis, legume plants can provide themselves and subsequent rotation crops with nitrogen, reducing requirements for environmentally and economically costly mineral nitrogen fertilization [2].

Root nodule symbiosis starts with a molecular dialogue between symbionts. The recognition of specific bacterial (Nod factors) and plant (flavonoids) signal molecules activates the symbiotic program controlling the formation of a nodule primordium and its colonization by rhizobia. Bacterial colonisation is provided by plant-derived infection threads. These tubular, cell wall-bound structures originate in root hairs and guide the growing bacterial colony towards the nodule primordium. There, rhizobia are released from the infection threads into the developing nodule cells. After their release, rhizobia multiply and develop into nitrogen-fixing bacteroids, gradually occupying host cells. Eventually, these processes result in the formation of mature root nodules, where nitrogen fixation occurs [3].

The past two decades of research on *Rhizobium*-legume interactions have provided unique insights into the genetics and molecular biology of nodulation. However, our understanding of symbiosis is largely restricted to the signalling necessary for its initiation and the development of dedicated organs [4]. By contrast, relatively little is known about the genetic mechanisms controlling or limiting the actual nitrogen fixation and symbiotic efficiency, which can be highly variable.

On the other hand, the available data indicate the important role of plant immunity in the *Rhizobium*-legume symbiosis [5]. Although plant defence pathways have not been well studied during symbiosis development, the transcriptomic data suggest that nodulation requires the fine regulation of plant immunity [6,7]. Plants have evolved diverse strategies to limit microbes through transcriptional activation of antimicrobial molecular mechanisms. It is conceivable that genes supporting such microbial control have been co-opted to control bacterial symbiosis. We hypothesized that negative regulators of symbiosis may be derived from ancestral defence genes through gene duplication and subsequent neo-functionalisation and can be identified by comparative phylogenetics. In turn, removal or modification of such negative regulators has the potential to enhance symbiotic performance in economically relevant legumes. Extensive genetic and transcriptomic resources of model legume *Medicago truncatula* [8,9] enable us to address this hypothesis.

Our survey of transcript levels of Medicago defence associated genes during the symbiosis with *Sinorhizobium meliloti* [8] identified the symbiosis-specific upregulation of the *β-Glucan-Binding Protein 1* (*GBP1*) gene. *GBP* genes encode dual domain proteins with glucan-binding and hydrolytic activities towards β-1,3/1,6-glucans [11,12,13]. The first GBP was isolated from soybean roots treated with *Phytophthora sojae* cell wall-derived glucans, elicitors of plant immune response. Later, several *GBP* genes were found in different legumes, including Medicago [14]. We have identified twelve members of the *GBP* family in the *Medicago truncatula* A17 r5.0 genome (Fig. 1A). During interactions with *S. meliloti, GBP1* showed the largest increases in expression relative to mock-inoculated control roots. By contrast to other *GBP* family members, *GBP1* is not upregulated upon challenge with either the pathogenic fungi *Botrytis cinerea* and *Rhizoctonia solani*, nor the oomycete *Phytophthora palmivora*, nor by treatment with laminarin (a branched glucan, structurally similar to glucans from cell walls of filamentous pathogens) (Fig. 1B-F).

**Fig. 1.**
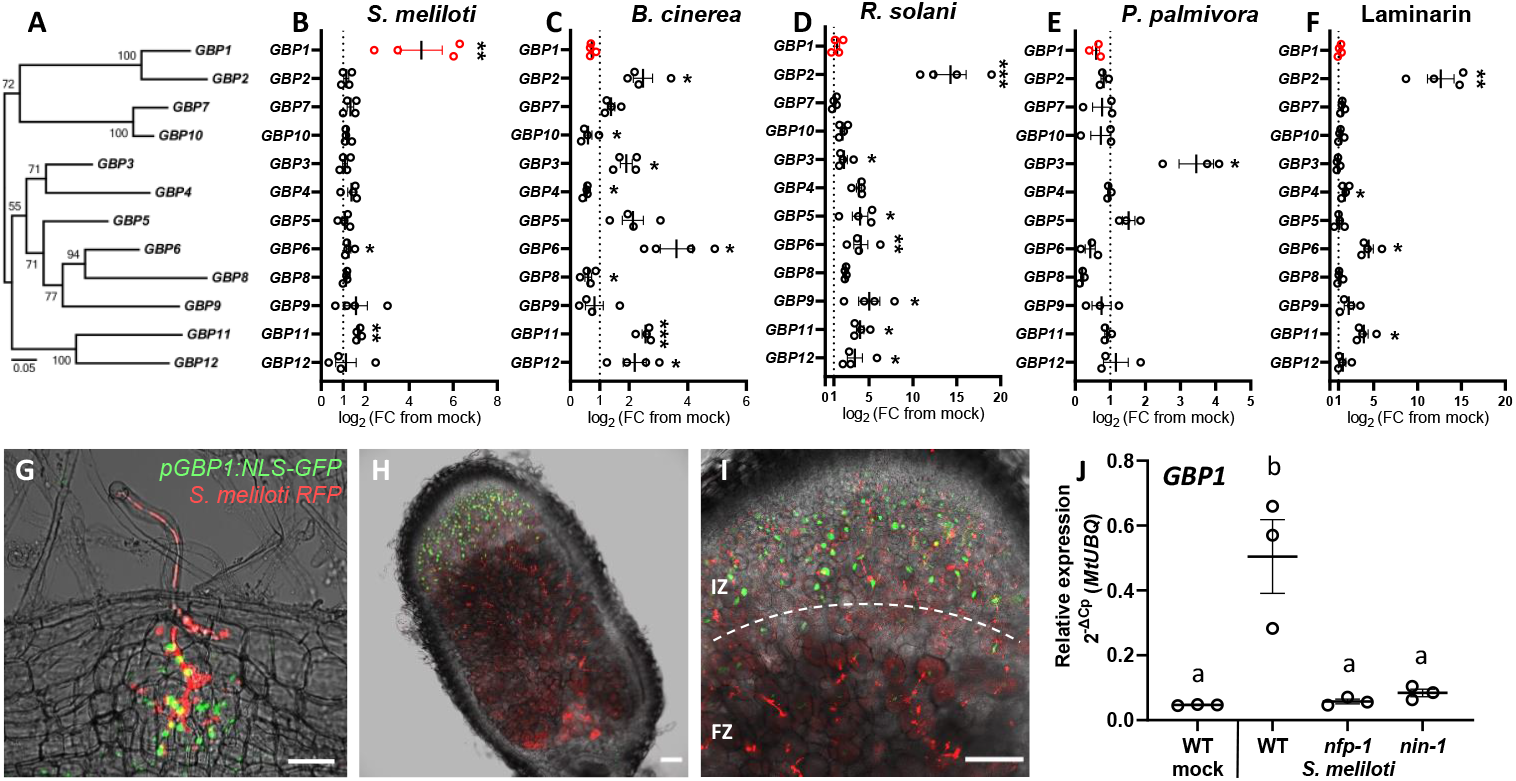
Expression of Medicago *GBP* genes and expression pattern of *GBP1* during symbiotic interactions with *Sinorhizobium meliloti*. (A) The Maximum Likelihood phylogenetic tree of Medicago GBP family proteins (number of bootstrap replications 1000; software MEGA X). Gene expression fold changes of *GBP* genes in roots colonized with *S. meliloti* (B), *Botrytis cinerea* (C), *Rhizoctonia solani* (D), *Phytophthora palmivora* (E) and laminarin treatment (F) compared to mock-inoculated roots (scatterplots show individual measurements; the horizontal bar represents the mean; error bars represent SEM; Student’s *t* test: \**P* < 0.05; \*\**P* < 0.01; \*\*\**P* < 0.001). Localization of *GBP1* gene expression assessed via a pGBP1:NLS-GFP fusion in root cortex (G) and nodules (H, F) colonized by *S. meliloti 2011* expressing RFP (IZ, infection zone; FZ, fixation zone; dashed line indicates the border between zones; scale bars: G, 50 µm; H, 200 µm; I, 50 µm). (J) Expression of *GBP1* gene in *nfp-1* and *nin-1* Medicago mutants (scatterplot shows individual measurements; the horizontal bar represents the mean; error bars represent SEM; one-way ANOVA with post hoc Tukey HSD test *P* < 0.05).

Using two promoter-reporter fusions (nucleus-targeted NLS-GFP and GUS-GFP), we evaluated the spatial expression patterns of *GBP1*. The *GBP1* promoter is activated during rhizobia infection of root hairs, root cortex, and nodule primordia (Fig.1G and Fig. S1A, B, C, E, F). In mature nodules, *GBP1* is predominantly expressed in the apex, with the strongest expression in the infection zone, gradually decreasing in the fixation zone (Fig.1H, I and Fig. S1D, G, H). The same expression pattern was detected by RT-qPCR analysis (Fig. S1I).

To investigate whether *GBP1* induction is a function of the symbiotic signalling and developmental pathways, we assessed *GBP1* transcript levels in *nfp-1* (*nod factor perception*) mutants which are defective in the perception of rhizobial Nod factors [15], and *nin-1* (*nodule inception*) mutants lacking a key transcription factor regulating symbiosis development [16]. Expression analysis shows that *GBP1* induction depends on Nod factor perception and NIN-mediated transcription regulation (Fig. 1J). This suggests that activation of *GBP1* during interactions with rhizobia is a part of the host symbiotic program.

To gain insight into the functional role of GBP1, we genetically deregulated *GBP1* expression using two approaches. First, we isolated Medicago *Tnt1* mutants [17] carrying a retrotransposon insertion in the *GBP1* promoter (*gbp1-1* and *gbp1-3*) or in the coding (*gbp1-4* and *gbp1-5*) sequences (Fig. S2A). *Tnt1* insertion into the *GBP1* promoter results in higher *GBP1* transcript levels, even in roots of mock-inoculated *gbp1-1* and *gbp1-3* plants. Line *gbp1-4* is a knockout mutant (Fig. S2B) and the insertion in the *gbp1-5* line is predicted to result in the overexpression of a truncated non-secreted GBP1 protein (OutCyte prediction [18], Fig. S2C-G).

Hypothesising that GBP1 could have a restricting effect on symbiosis, we assessed nitrogenase activity in nodules of *Tnt1* insertion mutants using an acetylene reduction assay. This is a robust method based on the ability of rhizobial nitrogenase to reduce acetylene to ethylene [19]. Remarkably, nodules of *gbp1-4* and *gbp1-5* mutants, lacking intact GBP1, showed higher nitrogenase activity compared to corresponding wild-type lines. Conversely, *GBP1*-overexpressing lines *gbp1-1* and *gbp1-3* produced nodules with decreased nitrogenase activity (Fig. 2A). Thus, plants lacking GBP1 are able to fix more nitrogen, which is supported by higher biomass accumulation of nodulated *gbp1-4* mutant under nitrogen-limited conditions. Vice versa, biomass accumulation of *gbp1-1* is decreased compared to the corresponding wild type (Fig. 2B). These findings suggest that GBP1 is a negative regulator of symbiotic nitrogen fixation and potentially should be regulated differently depending on the microsymbiont nitrogen-fixing capacity. To address this assumption, we compared interactions of Medicago cv. R108 with its optimal *S. meliloti* 2011 and inefficient symbionts *S. meliloti* 1021 and *S. medicae* WSM419 (Fig. S3A, B). Comparing these strains, we observed a strong correlation between the level of *GBP1* expression and the nitrogen-fixing capacity of rhizobia (r=0.9832; *P* < 0.0001; linear regression R^2^=0.9667; *P* < 0.0001) (Fig. 2C), suggesting a possible regulatory link between *GBP1* expression and microsymbiont efficiency.

**Fig. 2.**
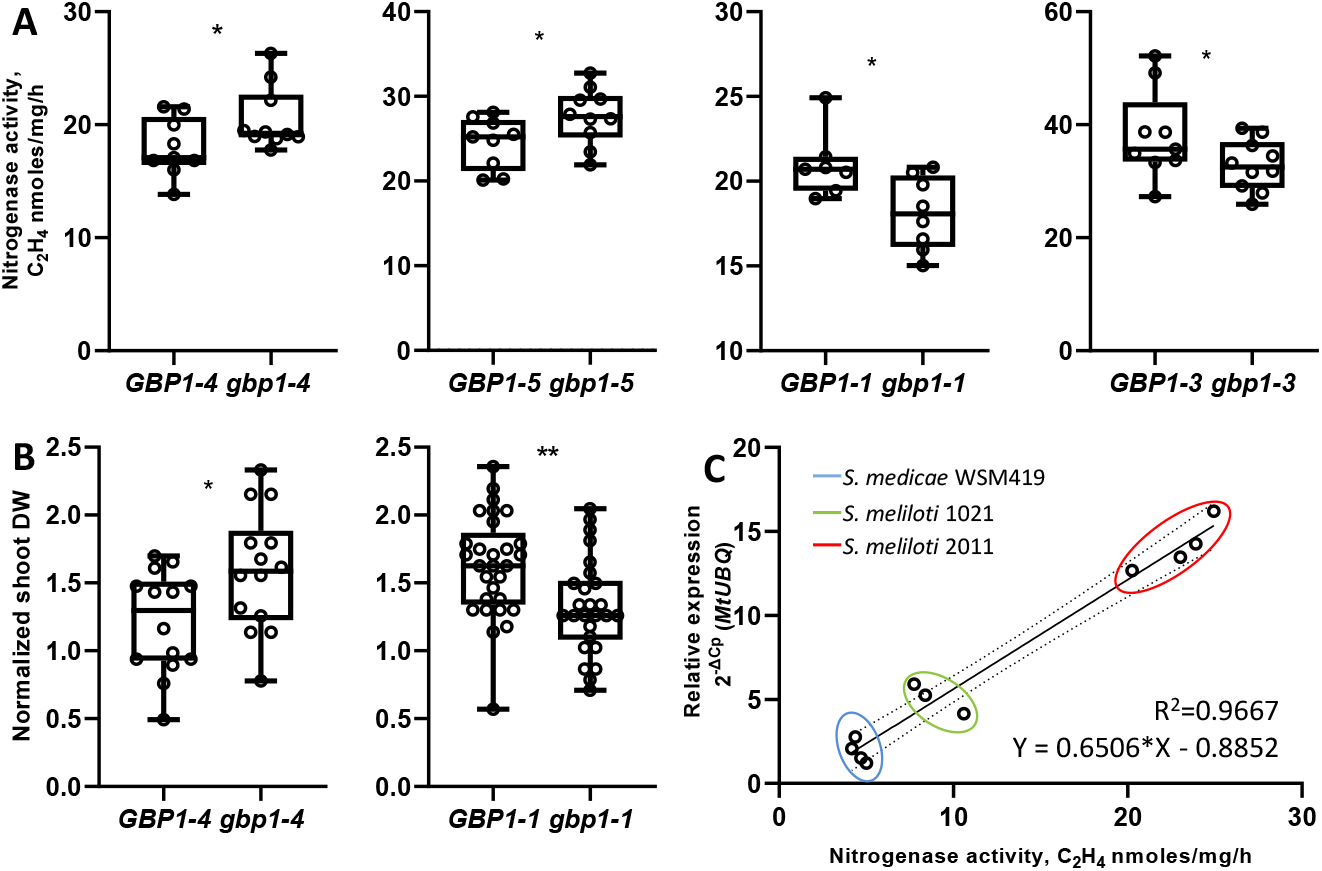
GBP1 is a negative regulator of symbiotic nitrogen fixation. (A) Nitrogenase activity in *GBP1* mutant and corresponding wild type nodules measured by the acetylene reduction assay. (B) Symbiotic biomass accumulation compared to mock-inoculated plants (box-and-whiskers plots show the lower and upper quartiles, the minimum and maximum values, the bar in the box represents the median, and the points represent individual measurements; Student’s *t* test: \**P* < 0.05; \*\**P* < 0.01). (C) Linear regression between *GBP1* expression level and nitrogenase activity in Medicago nodules established by three strains of rhizobia with different symbiotic efficiency (*P* < 0.0001).

Further mutant phenotyping showed that inactivation or truncation of GBP1 does not affect the level of nodulation. However, its upregulation in lines *gbp1-1* and *gbp1-3* significantly reduces the number of nodules per plant (Fig. 3A). None of the mutant lines were affected in their overall nodule development and morphology (Fig. 3B and Fig. S4A). To visualize nitrogen-fixing bacteroids in nodules, we used *S. meliloti* which express GFP under the *nifH* gene promoter. *NifH* encodes one of two main components of the nitrogenase complex [20]. Bacteroids in *gbp1-4* nodules had an elongated, rod-like shape similarly to wild type but were smaller in size. Strikingly, the GFP fluorescence intensity of these bacteroids was higher than wild-type nodule bacteroids (Fig. 3B, C, D), suggesting higher expression of the *NifH* gene. This is consistent with the higher level of nitrogenase activity of *gbp1-4* nodules. The smaller size of *gbp1-4* resident bacteroids indicates changes in their development, which occurs in the root nodule apex where *GBP1* is predominantly expressed. It possibly results in more bacteroids per infected plant cell, whose size remains as wild type (Fig. 3E).

**Fig. 3.**
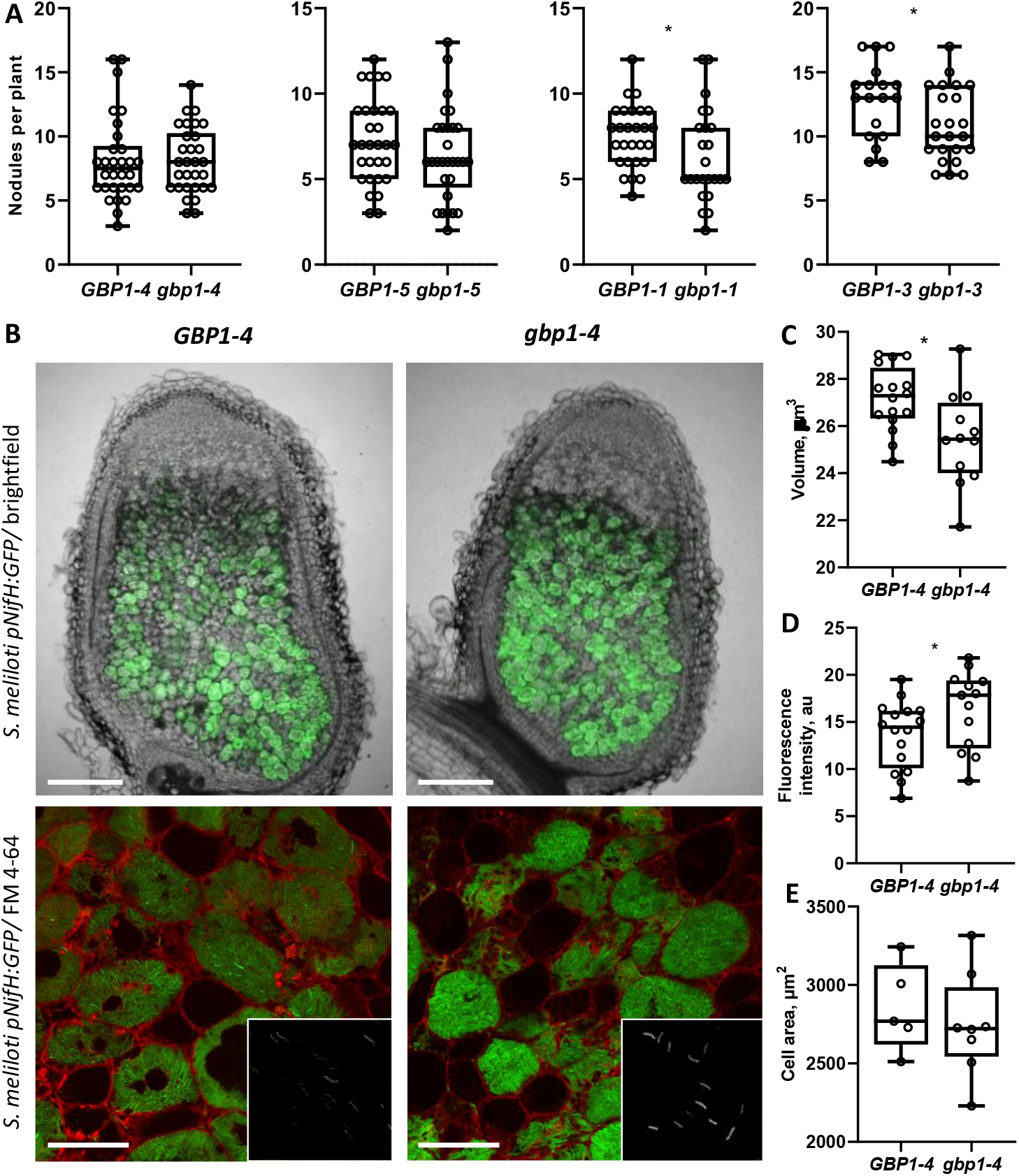
Root nodule phenotyping of *gbp1* mutants. (A) Quantification of nodulation of *gbp1* mutants and corresponding wild types. (B) Morphology of *gbp1-4* nodules at 15 days post inoculation with *S. meliloti* 2011 pNifH:GFP (scale bars: 250 µm; 50 µm). Quantification of the size (C) and fluorescence intensity (D) of bacteroids and the size of infected cells (E) form *gbp1-4* mutant line nodules and wild type nodules (box-and-whiskers plots show the lower and upper quartiles, the minimum and maximum values, the bar in the box represents the median, and the points represent individual measurements. Student’s *t* test: \**P* < 0.05).

In our second approach we genetically engineered Medicago roots to constitutively express *GBP1*. Composite plants overexpressing *GBP1* driven by strong ubiquitin (UBQ3) promoter were significantly impaired in their nodulation (Fig. S4B, C, D). Furthermore, the quantification of rhizobia colonizing rhizodermis of *pUBQ3:GBP1* expressing roots showed ∼2-fold decrease of bacteria compared to empty vector control (Fig. S4E), which can be an indication of potential antimicrobial properties of GBP1.

*GBP* genes are widespread among land plants; however, this family is particularly enlarged in legumes. Most of the analysed diploid dicot and monocot plants have up to three *GBP* genes, whereas in diploid legumes this amount ranges from six (*Lotus japonicus, Cajanus cajan, Lupinus angustifolius*) to twelve in Medicago (Fig. S5). Given their expanded family size across legumes, we looked for evidence of *GBP1* functionality in economically relevant legumes. Our phylogenetic analysis shows that red clover (*Trifolium pratense*), pea (*Pisum sativum*), faba bean (*Vicia faba*), soybean (*Glycine max*), common bean (*Phaseolus vulgaris*), cowpea (*Vigna unguiculata*) and pigeon pea (*Cajanus cajan*) share *GBP* genes in the same subclade and hence might have the same functionality. Like Medicago *GBP1*, some of these genes show transcriptional upregulation in response to rhizobial colonization of pea and soybean [21, 22, 23]. These data provide initial evidence that symbiosis-inducible GBP genes may quantitatively modulate nitrogen fixation in grain legumes.

*GBP1* is a unique example when knockout of the symbiotically induced gene increases nitrogenase activity, resulting in higher biomass production. Our findings offer new strategies to sustainably improve nitrogen nutrition through mutagenesis breeding or targeted genome modification in crop legumes.

## Material and Methods

### Plant materials

*M. truncatula* cv R108 and cv Jemalong A17 were used in this study. Mutants harbouring *Tnt1* insertions within *MtGBP1* coding and regulatory sequences (*gbp1-1*, NF19905_high_1; *gbp1-3*, NF16527_high_3; *gbp1-4*, NF1807-Insertion-17; *gbp1-5*, NF15880_high 1) were identified in a *Tnt1*-insertion mutant population of *Medicago truncatula* by sequencing the PCR-amplification products. The relevant primers are shown in Supplemental Table 1. Homozygous mutant and corresponding wild type lines were isolated from self-pollinated heterozygous individuals. The line carrying the *gbp1-4* insertion was backcrossed to R108 wild type and resegregated. Homozygous *GBP1-4* progeny of the same parent was isolated and used in subsequent experiments as a wild type control.

Medicago seeds were surface sterilized by incubation for 5 min in concentrated sulphuric acid, followed by six washes in sterile water, then 5 min in 10% hypochlorite (commercial bleach), followed by seven washes in sterile water. Sterilised seeds were plated on wet filter paper, vernalized for 2 days at 4 °C and germinated at 25 °C for 24 h in darkness.

### Nodulation

One day old seedlings were sown on a 1:1:1 mix of vermiculite, Terragreen and perlite saturated with Färhaeus medium (1 mM MgSO4·7H2O, 0.75 mM KH_2_PO_4_, 1 mM Na_2_HPO_4_, 15 µM Fe‐citrate, 0.75 mM Ca(NO_3_)_2_, 0.7 mM CaCl_2_, 0.35 µM CuSO_4_·5H_2_O, 4.69 µM MnSO_4_·7H_2_O, 8.46 µM ZnSO_4_·7H_2_O, 51.3 µM H_3_BO_3_, 4.11 µM Na_2_MoO4·2H_2_O, pH 6.7) and grown in a growth chamber at 21 °C and 16/8-h light/darkness. Three days after germination plants were inoculated with *Sinorhizobium meliloti* 2011, *Sinorhizobium meliloti* 1021 or *Sinorhizobium medicae* WSM419 [10] (OD600=0.1, 2 mL per plant). Nodulated roots were collected for analysis 21 days after inoculation. Plant dry weight was measured and normalized to water inoculated plants (mock control).

### Pathogen infection and laminarin treatment

Bleach-sterilised seeds of *Medicago truncatula* were germinated and transferred on sterile plates with 0.8% agarose. For *P. palmivora* infection assays, two-day old seedlings were inoculated with 10 μl of *P. palmivora* LILI-td [24] zoospore suspensions (5 × 10^4^ zoospores/ml). Twenty-four hours after inoculation, infected roots were pooled into four biological samples for RNA extraction. For *B. cinerea* R5 infection assays, five-day old seedlings were inoculated with 100 μl of *B. cinerea* spore suspensions (5 × 10^4^ conidia/ml). Two days after inoculation, infected roots were pooled into four biological samples for RNA extraction. For *R. solani* infection assays, square plastic Petri dishes (120mm × 120mm) with five-day old seedlings grown on agar plates were inoculated with four mycelial plugs (5 mm × 5mm) taken from the leading edge of one-week old, inoculated potato dextrose agar Petri dish cultures. Twenty-four hours after inoculation, infected roots were pooled into four biological samples for RNA extraction. For laminarin treatment, four dayold seedlings were treated individually with 100 µl of 4 µg/ml solution of laminarin (MERCK, L9634). After two hours of treatment, roots were pooled into four biological samples for RNA extraction.

### Gene expression analysis

RNA was extracted using the RNeasy Mini Kit including on-column DNAse digest according to manufacturer recommendations (Qiagen). Reverse transcription and cDNA synthesis were performed on 1 µg of total RNA using the iScript cDNA Kit according to manufacturer recommendations (Bio-Rad). Quantitative PCR (qPCR) was performed in technical triplicates using SYBR Green I Master kit in a LightCycler® 480 (Roche). Ten microliter reaction volumes were used with 7.5 µl of master mix containing 1 µM gene-specific primers and 2.5 µl of 10-fold pre-diluted cDNA.

### Acetylene reduction assay

Nitrogenase activity was measured by the acetylene reduction assay [25]. Nodulated roots were collected into 13-ml tubes. Tubes were stoppered with rubber septa (Suba-Seal n°29) and each injected with 1 mL of acetylene. After one hour of incubation, the emitted ethylene gas was quantified using a Perkin Elmer Clarus 480 gas chromatograph equipped with a HayeSep N (80–100 MESH) column. The injector and oven temperatures were kept at 100°C, while the FID detector was set at 150°C. The nitrogen carrier gas flow was set at 8–10 mL/min. Nitrogenase activity is reported as nmol of ethylene/mg nodules/hour.

### Promoter GUS assay

Transformed *M. truncatula* roots were collected and washed twice in 0.1 M sodium phosphate buffer, pH 7.2, incubated in GUS buffer (100 mM sodium phosphate pH 7.0, 5 mM K_3_Fe(CN)_6_, 5 mM K_4_Fe(CN)_6_ and 2 mM 5-bromo-4-chloro-3-indoxyl-b-d-glucuronic acid, X-gluc) under vacuum at room temperature for 30 min to allow the buffer to replace air in the tissue, incubated at 37°C for 2 h to enable the enzymatic reaction, and analyzed using Leica M165 FC and Zeiss Axioimager M2 microscopes.

### Microscopy

The selection of transgenic roots and nodules was performed on a Fluorescent Stereo Microscope Leica M165 FC equipped with a DFC310FX camera and DSR filter (10447412) to detect DsRED reporter and GFP2 (10447407) to detect GFP expressing rhizobia. Confocal imaging of GFP-fused proteins was done on transgenic hand-sectioned roots or nodules by using a Leica TCS SP8 confocal microscope with emission/excitation settings 488/507 nm for GFP, 587/610 nm for RFP, 515/640 nm for FM4-64 (working concentration 30 μg/mL; nodule sections were stained for 1h on ice). The software Imaris and ImageJ were used for image processing and quantitative analysis.

### Phylogenetic analysis

In total, 98 protein sequences were retrieved from GenBank and via BLAST using the respective *M. truncatula* GBP proteins. Sequences were aligned using MAFFT v7.313 [26] and subjected to Maximum Likelihood analysis with IQTREE v1.6.7 [27]. The tree was visualized and annotated through iTOL v5.6.3 [28]. Branches corresponding to partitions that were reproduced in less than 50% of bootstrap replicates were collapsed.

For Medicago GBP protein family, sequences were aligned using CLUSTAL W alignment, and phylogenetic analysis was performed in MEGA 11 [29], using the Maximum Likelihood method, and displayed in a bootstrap consensus tree inferred from 500 replicates.

## Acknowledgments

We would like to thank Rene Geurts (Wageningen University) for providing a fluorescently labelled strain of *S. meliloti*, Uta Paszkowski (University of Cambridge) for *R. solani;* and Phil Poole (University of Oxford) for constructive discussions.

## Author contributions

A.G. designed and conducted the experiments. M.B.B., E.E., R.D. conducted the experiments. A.G. and S.S. wrote the manuscript.

## Data and materials availability

All materials are available upon request.

---

**Fig. S1.**
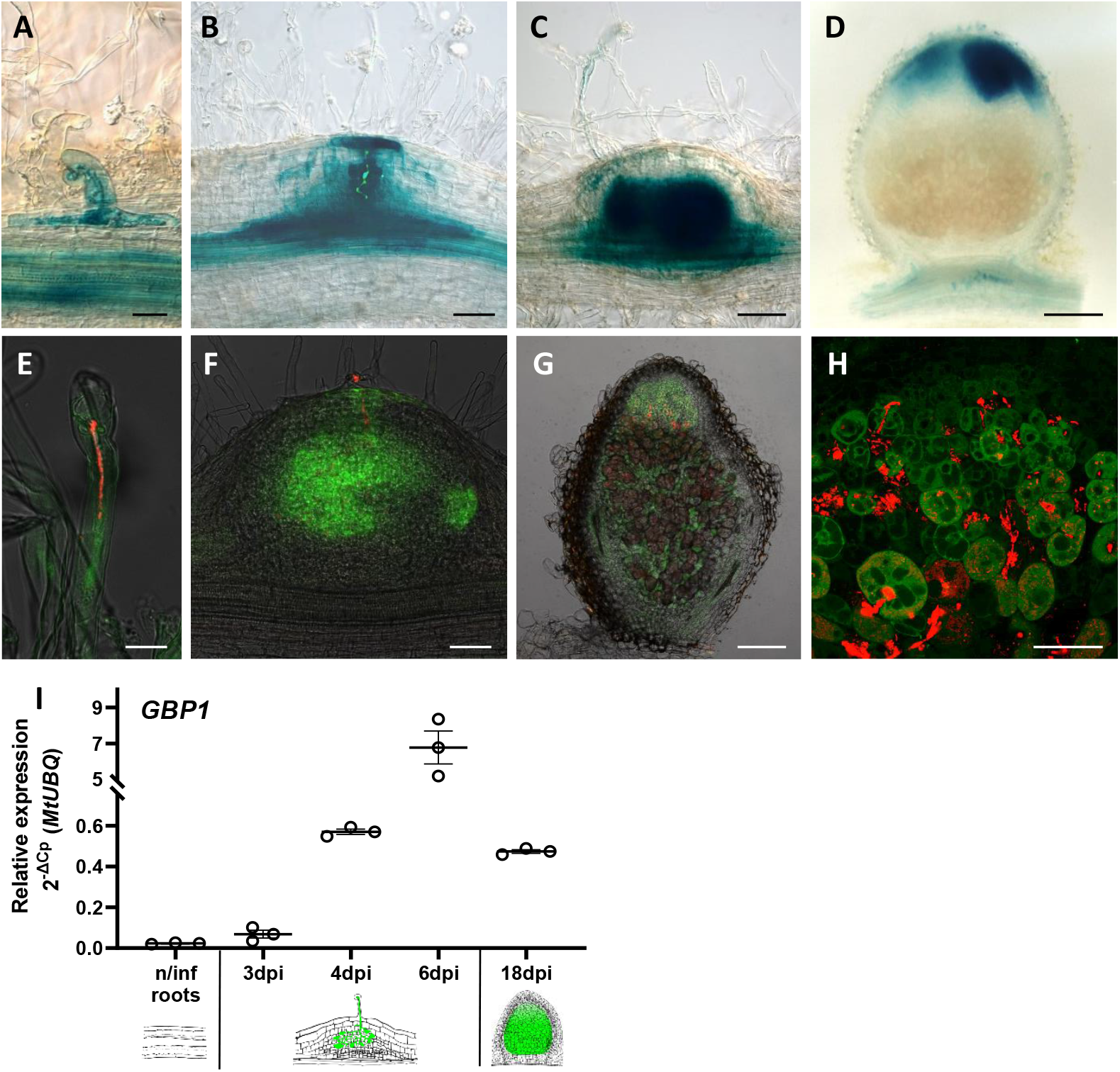
Expression pattern of Medicago *GBP1* gene during root nodule development. Localization of *GBP1* gene expression assessed using pGBP1:GUS-GFP fusion in root hairs (A, E), root cortex (B), nodule primordia (C, F) and nodules (D, G, H) colonized by *S. meliloti 2011* expressing RFP (scale bars: A, E, 20 µm; B, C, D, G 200 µm; F, 100 µm; H, 50 µm). (I) Transcript level of GBP1 in Medicago roots colonised by *S. meliloti* 2011 at different time points (dpi, days post inoculation; scatterplot shows individual measurements; the horizontal bar represents the mean; error bars represent SEM).

**Fig. S2.**
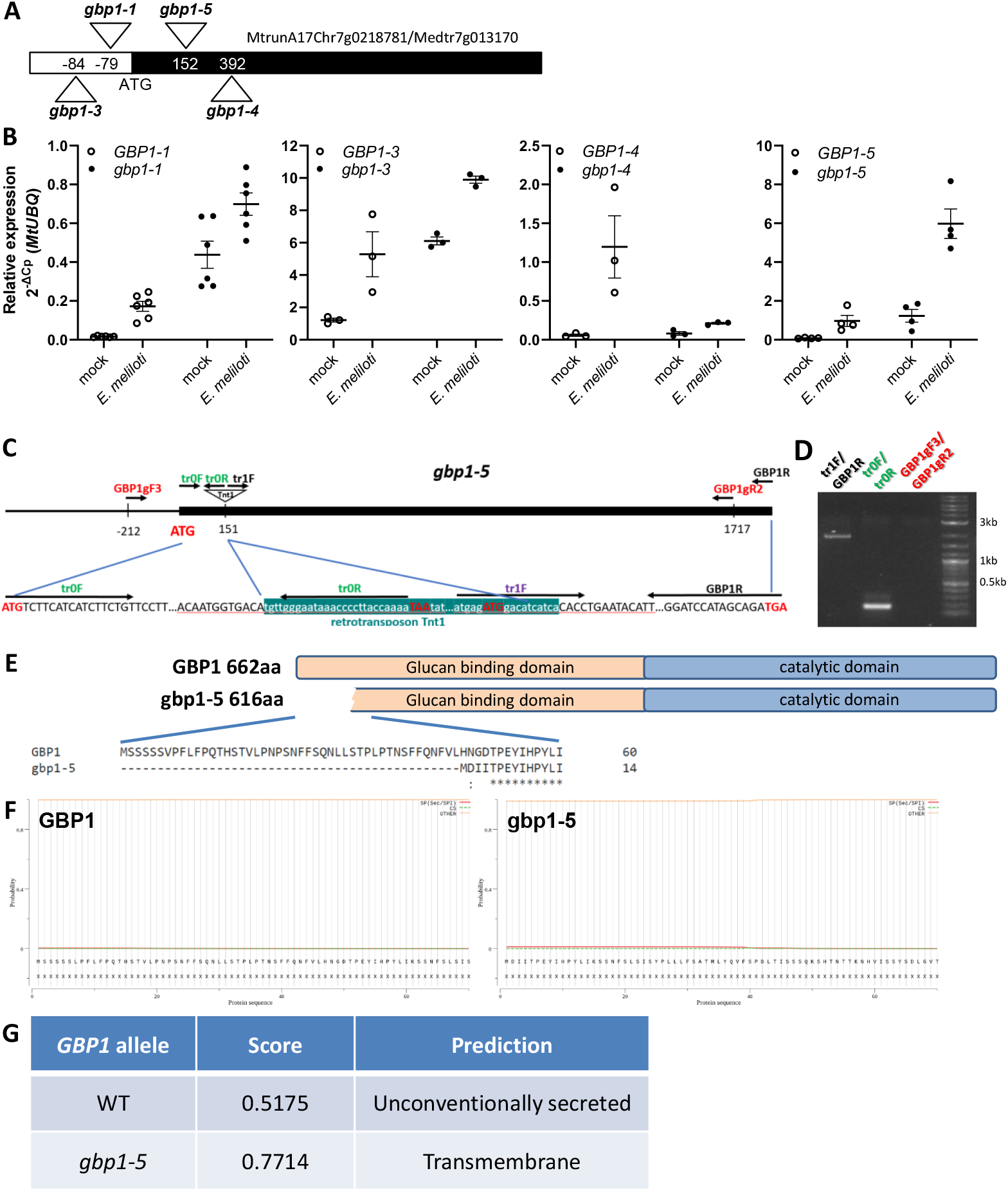
Isolation and characterization of Medicago *gbp1* mutants. (A) Gene model showing the positions of Tnt1 retrotransposon insertion sites in the *GBP1* gene (MtrunA17Chr7g0218781/Medtr7g013170). (B) RT-qPCR performed on RNA extracted from nodulated roots of *gbp1* mutants. (C) Gene model showing the positions of *gbp1-5* retrotransposon insertion site splitting the gene transcript. (D) RT-PCR performed on RNA extracted from nodulated roots of *gbp1-5* mutant to detect two resulting transcripts. Primer positions are shown in (C). (E) Protein model of truncated *gbp1-5* missing 46 amino acids. (F) Signal peptide prediction for GBP1 (wild type) and truncated protein (*gbp1-5*). (G) OutCyte prediction of unconventional protein secretion (without a signal peptide) for GBP1 wild type and gbp1-5 truncated proteins.

**Fig. S3.**
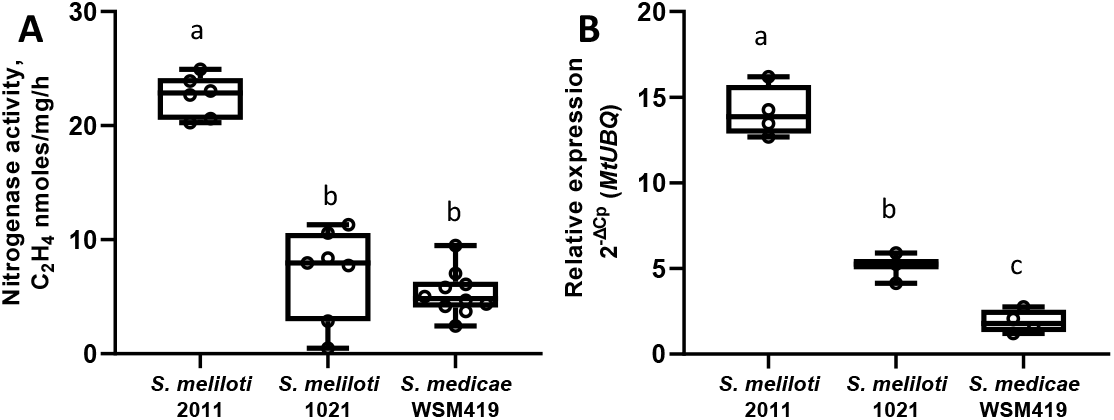
Comparative study of interactions of Medicago cv. R108 with its optimal *S. meliloti* 2011 and inefficient symbionts *S. meliloti* 1021 and *S. medicae* WSM419. (A) Nitrogenase activity in nodules initiated by different rhizobia measured by the acetylene reduction assay. (B) Expression level of *GBP1* gene in nodules initiated by different rhizobia (box-and-whiskers plots show the lower and upper quartiles, the minimum and maximum values, the bar in the box represents the median, and the points represent individual measurements; one-way ANOVA with post hoc Tukey HSD test *P* < 0.05).

**Fig. S4.**
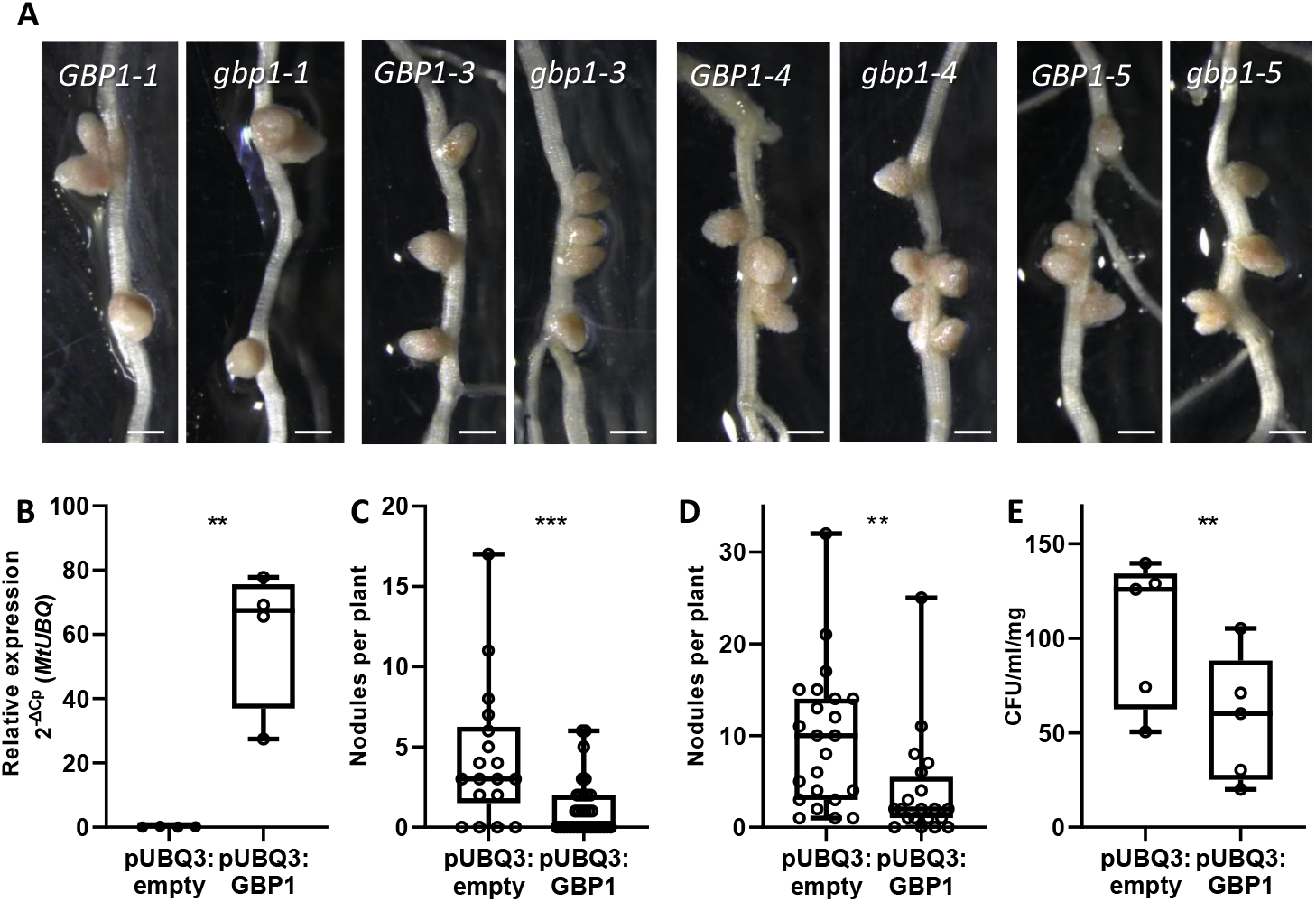
Genetic deregulation of *GBP1* expression. (A) Nodule developed on root of *gbp1* mutants (scale bars 1mm). (B) Expression level of *GBP1* gene in roots expressing pUBQ3:GBP1 and empty vector control. Level of nodulation of roots expressing pUBQ3:GBP1 and empty vector control at 10 dpi (C) and 17 dpi (D). (E) Quantification of rhizobia colonizing rhizodermis roots expressing pUBQ3:GBP1 and empty vector control (box-and-whiskers plots show the lower and upper quartiles, the minimum and maximum values, the bar in the box represents the median, and the points represent individual measurements; Student’s *t* test: \**P* < 0.05; \*\**P* < 0.01).

**Fig. S5.**
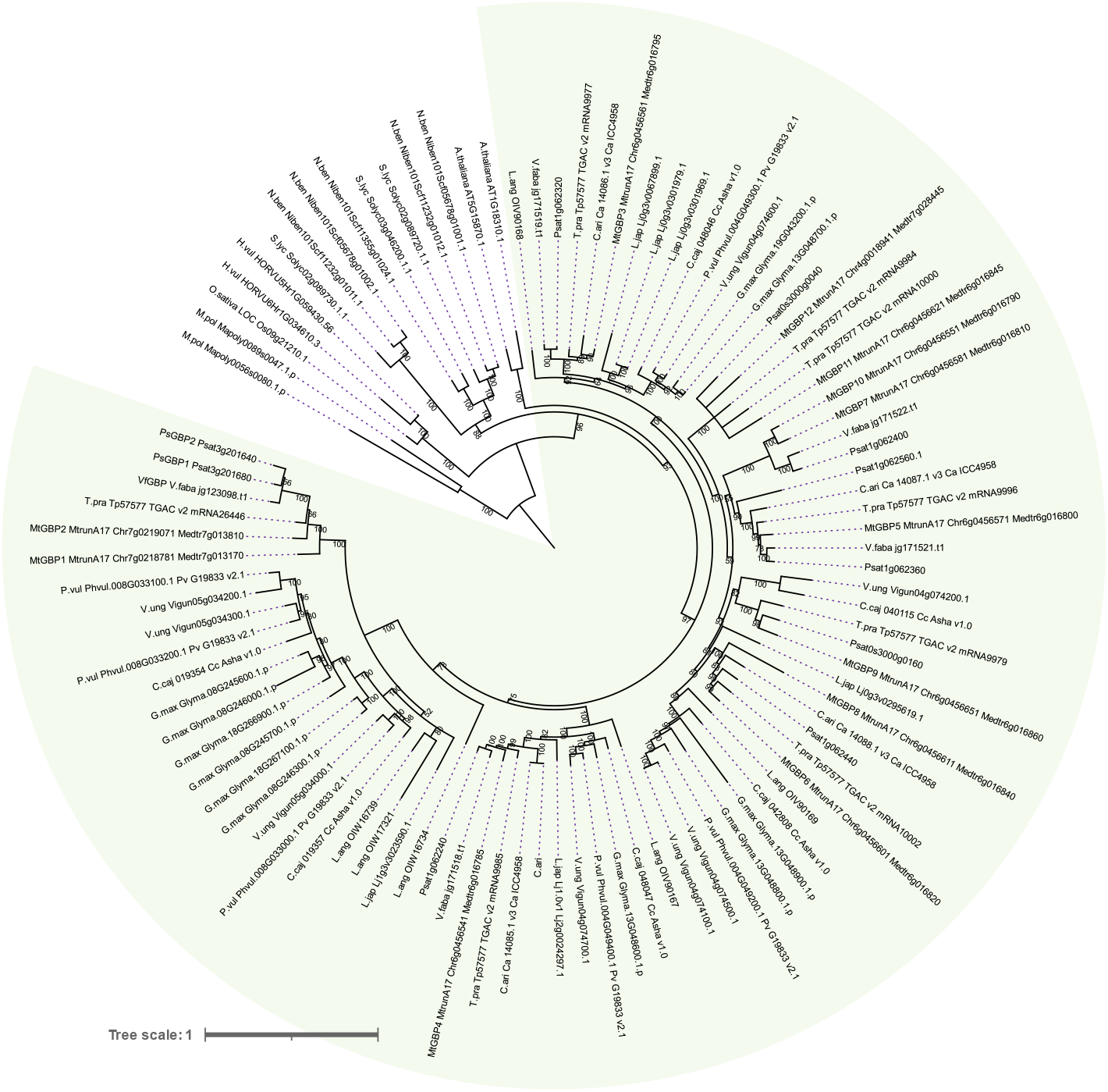
Maximum likelihood tree of GBP1 family reconstructed with IQ-TREE on an alignment of 98 protein sequences from 17 species. Legumes are highlighted in green.

